# Widespread arginine phosphorylation in human cells - a novel protein PTM revealed by mass spectrometry

**DOI:** 10.1101/725291

**Authors:** Songsen Fu, Chuan Fu, Quan Zhou, Rongcan Lin, Han Ouyang, Minning Wang, Ying Sun, Yan Liu, Yufen Zhao

## Abstract

Arginine phosphorylation (pArg) is recently discovered as a ubiquitous protein N-phosphorylation in bacteria. However, its prevalence and roles in mammalian cells remain largely unknown due to the lack of established workflow and the inherent lability of the phosphoramidate (P-N) bond. Emerging evidence suggests that N-phosphorylation may extensively exist in eukaryotes and play crucial roles. We report an experimental phosphoproteomic workflow, which for the first time allowed to reveal the widespread occurrence of pArg in human cells by mass spectrometry. By virtue of this approach, we identified 152 high-confidence pArg sit]es derived from 118 proteins. Remarkably, the discovered phosphorylation motif and gene ontology of pArg hint a possible cellular function of arginine phosphorylation by regulating the favorability of propeptide convertase substrate. The generated extensive data set should enable a better understanding of the biological functions of eukaryotic pArg in the future.

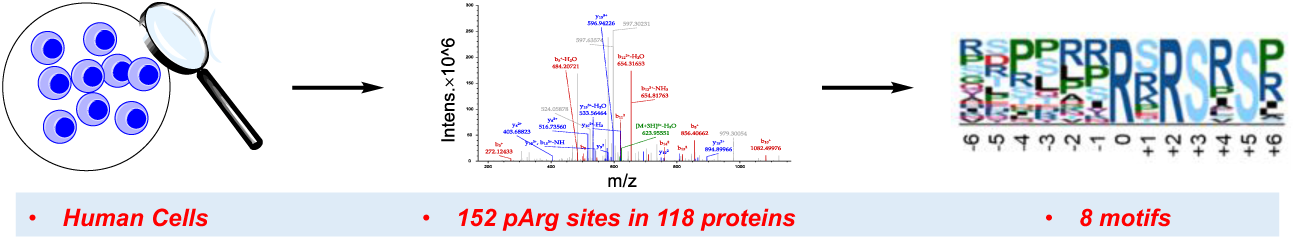

## INTRODUCTION

Protein reversible phosphorylation is a ubiquitous and dynamic post-translational modification (PTM) involved in virtually all aspects of cellular regulation.^1-3^ For decades, extensive studies were focused on *O*-phosphorylation taking place on serine, threonine and tyrosine residues, while *N*-phosphorylation of arginine, histidine and lysine has long been underestimated due to the intrinsic instability of phosphoramidate.^4, 5^ Lagging behind the striking discoveries of histidine phosphorylation in mammalian cells,^6^ the study of arginine phosphorylation (pArg) has just begun. Recent breakthrough studies shed light on pArg participating in DNA transcription regulation and protein degradation in gram-positive bacteria.^7-9^ The presence of pArg in vertebrates is sporadically reported, such as on histone H3,^10, 11^ histone H4,^12^ and myelin basic protein (MBP).^13^ However, its prevalence and roles in eukaryotes remain largely unknown, mainly due to the limited tools and the lack of appropriate experimental workflow.

The large-scale discovery of pArg in mammalian cells has been hindered by several issues. Firstly, the intrinsic nature of P-N bond in pArg makes it unstable to heat and acid.^14^ Consequently, the commonly used proteomic protocols designed for *O*-phosphorylation, involving the use of strong acids, are not suitable. Secondly, sufficient amount of identified pArg proteins is necessary for bioinformatic analysis. Due to mammalian pArg kinase and phosphatase has not yet been discovered, the commonly used strategies for prokaryotes, such as heat-shock treatment and knock-out of the corresponding phosphatase to elevate pArg level, are not applicable.^14^ Thirdly, as a PTM, pArg level may vary in different cells. Unfortunately, there is no precedent for choosing the appropriate mammalian cell lines.

Herein, we reported the large-scale discovery of pArg in human cells by a new analytical workflow. Crucially, 118 high-confidence pArg proteins fell into several specific protein classes participated in DNA/RNA associated process. The pArg motif was also deduced which paves the way for the identification of pArg kinase and phosphatase in the future. Strikingly, a pArg site was found in the vital cleavage region of proglucagon (PG), suggesting a mechanism of how pArg modulates the substrate favorability of prohormone convertase (PC). The systematic discovery of human pArg proteins not only clarified the long-standing question of its occurrence in eukaryotes but also provided insights into its biological studies in the future.

## RESULTS AND DISCUSSION

### pArg level exhibits variability in different cell lines

To choose a suitable model cell line, we first analyzed the total pArg content in human cell lines with the assistance of home-made pan-specific pArg antibody.^15^ The specificity of our antibody and its applicability for complex samples were validated using cell lysate of *Bacillus subtilis* and *Bacillus licheniformis* (Fig. S1). Subsequently, we applied Western blot assay to the human cell lysate and found pArg showed a differential profile across cell lines (Fig. S2).For Jurkat cell line, several high-intensity bands were significantly reduced or eliminated after acid-treatment (Fig. 1), further demonstrating that detected signals were acid-labile pArg. In consideration of signal intensity and protein diversity, we decided to choose Jurkat cells as a model for proteomics study.

**Figure 1.**
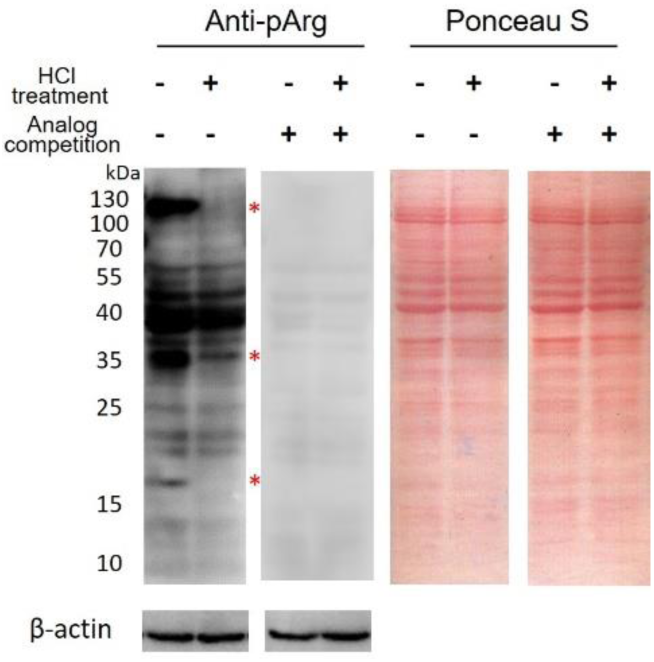
Western blot analysis of Jurkat cell lysate by pan-specific pArg antibody and Ponceau staining. As a control, the lysate was treated with 0.2 N HCl at 30 °C for 1 hr or blotted with an analog-competition assay. Bands with a markedly weakened intensity after HCl-treatment are indicated by red asterisks. The blot was slightly overexposed to show the difference clearly. The purpose of acid-treatment is mainly to show the difference before and after the treatment. However, prolonged acid-treatment was avoided which can precipitate eukaryotic proteins from the lysate for an unknown reason,

### Optimization of sample preparation and precursor fragmentation mode substantially increases the number of identified pArg sites

Encouraged by the direct detection of pArg proteins by Western blotting, we set out to explore the prevalence of pArg in Jurkat cells using mass spectrometry-based proteomics. In general, the large-scale identification and precise localization of labile *N*-phosphorylation sites greatly rely on suitable sample preparation approach, proper chromatographic separation, and appropriate precursor fragmentation mode.^16^ Also, the enrichment of phosphopeptide is crucial for all substoichiometric PTMs. However, an insurmountable obstacle was the acid lability of pArg, since the acid condition was commonly considered as a necessity for above-mentioned requirements. Moreover, sequence discrimination arose by enrichment also needs to be considered, because any serious deviation from the physiological distribution will affect deducing pArg motif. Accordingly, we adopted the reported mild acidic TiO_2_-based enrichment method^14^ as a blueprint and designed an improved workflow (detailed in Table S1).

The entire proteomic workflow was optimized in several aspects to expand the coverage of phosphopeptides while preserving labile pArg (Fig. 2a). These included 1) treating cells with a broad-spectrum phosphatase inhibitor in prior to cell lysis, 2) inactivating endogenous phosphatase with a strong denaturant, 3) removal of small-molecule interferences by acetone precipitation, 4) control of environmental temperature and shortening sample processing time, 5) improving the speed of TiO_2_ enrichment, 6) using volatile eluent to avoid extra off- or on-line desalting steps, 7) screening MS fragmentation mode and relevant parameters.

**Figure 2.**
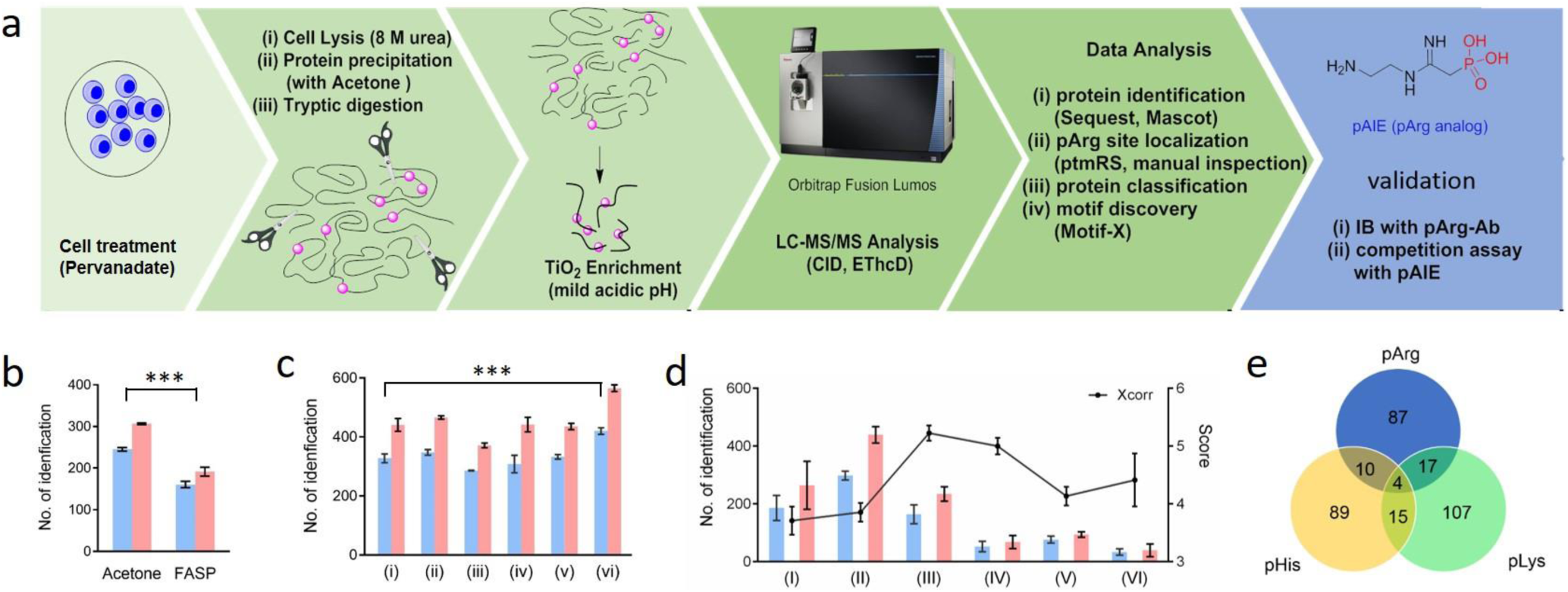
Experimental workflow and summary of results. (**a**) Cells are pre-treated with sodium per-vanadate. After cell lysis with 8 M urea, protein precipitation by acetone, and tryptic digestion, phosphopeptides are enriched with TiO_2_ under weak acidic condition and analyzed by LC-MS/MS. Proteins of interest are then biochemically verified. The proteomics workflow is optimized in the aspects of (**b**) sample preparation, (**c**) cell pre-treatment and (**d**) MS^2^ fragmentation mode. The number of identified pArg protein and peptides are indicated with blue and red bars, respectively. Centre values represent the mean; error bars indicate the s.d. for *n*=2 technical replicates. The conditions of (i)∼(vi) and (I)∼(VI) are detailed in Table S2 & S3. Statistical analysis is performed with Graphpad Prism 8, and the significance (p<0.05) is indicated with ***. (**e**) The Venn diagrams of high-confidence pArg, pLys, and pHis proteins.

As a result, these optimizations substantially increased the number of identified pArg sites. As shown in Fig. 2b, acetone precipitation following Urea-based cell lysis increased the number of identified pArg proteins and peptides from 161 and 192 to 245 and 307, as compared to the standard FASP method.^17^ Pre-treatment of cells with sodium per-vanadate (condition vi) further boosted identities to 420 and 565 (Fig. 2c). Presumably, per-vanadate serving as a covalent phosphatase inhibitor is more potent than Na_3_VO_4_ or pAIE (competitive inhibitor). Furthermore, we compared three MS^2^ fragmentation modes and six parameters setting (Table S2), including collision-induced dissociation (CID) (condition II), the combination of electron-transfer dissociation (ETD) and higher-energy CID (EThcD, III, IV & V), and neural loss triggered EThcD (I & VI). As shown in Fig. 2d, condition I, II and III produced the greatest number of identification and the highest cross-correlation score (Xcorr) in the respective mode. Accordingly, two biological replicates from condition I, II, and III, respectively, were combined as a “union dataset”.

We finally identified 817 pArg sites of 730 peptides derived from 500 proteins in Jurkat cells, demonstrating the widespread occurrence of pArg in human, a representative of senior mammals. Meanwhile, the unbiased TiO_2_-based enrichment allowed us to analyze *O*- and *N*- phosphorylation species simultaneously. In general, the per-vanadate treatment improved the identification of all six phosphorylation forms (Fig. S3). Phosphopeptides accounted for about 2/3 of eluted peptides, indicating good selectivity of enrichment procedure (Fig. S3, Excel S1). Of note, pArg proteins only showed limited over-lapping with pHis and pLys (Fig. 2e). The *ptm*RS distribution of pArg was similar to that of pHis and pLys (Fig. S4), but the proportion of high-confidence category (*ptm*RS ≥ 75) was lower than that of *O*-phosphorylation (Fig. S5). This was presumably due to the fact that phosphoramidate is more susceptible to neutral loss than phosphoester in the gas-phase.^18, 19^ Despite the use of *ptm*RS > 99 as a standard for reliable site localization of *O*-phosphorylation site, we relaxed this standard to 75%, which is used for *N*-phosphorylation in the literature.^16, 18^ Although this operation sacrificed the false localization rate (FLR) to some extent, it will result in a significant increase in identity, increasing the FLR to about 1% at an acceptable cost.^20, 21^ In order to verify the credibility of our data processing approach, we searched a recently published *E.coli* dataset reporting 246 pHis sites.^16^ Since pArg is absent in Gram-negative bacteria, *E.coli* can be used as a negative control to evaluate the parameters of data analysis. As a result, we only identified two ambiguous pArg sites in *E.coli* dataset with above-mentioned filter. In addition, the total number and relative ratio of *O*-phosphorylated proteins identified in the Jurkat sample were comparable with previous studies on the human phosphoproteome,^22, 23^ and there was only limited overlap between three *N*-phosphorylation forms. Taken together, the 152 high-confidence pArg sites identified by our new workflow were statistically reliable (Excel S2).

### Human pArg protein enriches in specific protein classes and shows preferable phosphorylation motifs

Gene ontology (GO) analysis was performed to elaborate on the association of pArg with specific physiological functions in human cells. Briefly, the “union dataset” was searched with SEQUEST, and then peptide hits with a C-terminal pArg site were excluded because they are less likely to be cleavable by trypsin.^26^ After manually checking the quality of spectrum, 118 pArg proteins were used for GO analysis. Protein classification revealed that pArg proteins were strongly enriched in the group of “nucleic acid binding”, together with a substantial number of “transcription factor”, “hydrolase”, “enzyme modulator”, “chaperone” and “membrane traffic protein” (Fig. 3a). Strikingly, the first and the only well-studied prokaryotic pArg protein CtsR is also a transcription repressor regulating gene expression through binding to DNA. Upon phosphorylation on Arg residues, CtsR was impaired by converting the positive-charged DNA binding domain to negative. Although the same pArg “switch” may exist in eukaryotic transcriptional regulation, to our knowledge, our findings are the first evidence supporting this hypothesis.

**Figure 3.**
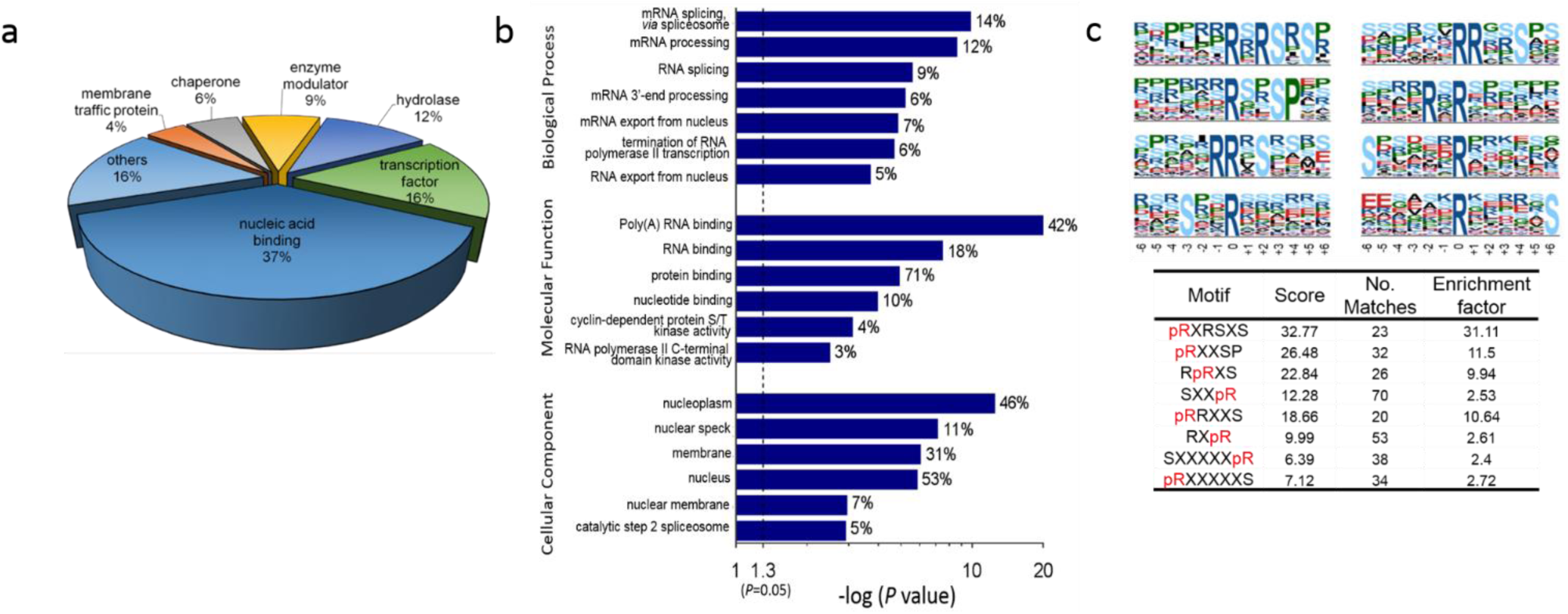
Bioinformatic analysis of the high-confidence pArg proteins and sites. (**a**) pArg proteins classification obtained by using PANTHER algorithm. Only categories of >3 members were used for analysis. (**b**) GO analysis of pArg proteins with DAVID software in terms of biological process, molecular function and cellular component. The statistical significance is plotted on the x-axis as –log (*p*-value); the percentage (%) of each category is indicated on the right of the bar. The dashed line represents the threshold of 1.3 (*p*-value = 0.05). (**c**) pArg motifs revealed with Motif-X algorithm.^24, 25^ The significance threshold of motif assignment is set to *p* < 10^−6^, and the minimum number of matches is set to 20.

Further bioinformatics analysis pinpointed that pArg proteins in Jurkat cells predominantly correlated with “mRNA associated process”, such as “mRNA splicing and processing”, “mRNA exporting from nucleus” and “termination of RNA polymerase II transcription” (Fig. 3b, Fig. S6, Excel S3). This result differs from that observed in Gram-positive bacteria, where pArg links with protein synthesis/degradation and metabolic functions.^9, 14^ Furthermore, we found that the cellular localization of human pArg proteins corresponds to their biological processes, showing enrichment in the nucleus. Interestingly, pArg exhibited some similarities to pLys in terms of biological process and cellular component (Fig. S7-S10). Compared to pHis and pLys, pArg sites were exclusively present in certain protein kinases, implying its potential biological functions associated with kinases.

The pHis proteins cellular location (in both nucleoplasm and cytosol) obtained by our study, if serving as a positive control, was consistent with the result previously obtained by immune-fluorescence staining on Hela cells.^6, 27^ Such consistency further demonstrated that the cellular localization and function of pArg proteins revealed by our study is credible.

Importantly, human pArg possessed consensus phosphorylation motifs which were not observed in prokaryotes. The statistical analysis on 141 high-confidence pArg sequences yielded 8 degenerated sequons (Fig. 3c), and four of them have an enrichment factor greater than 10, relative to human proteome as a background. In particular, Ser and Pro were high-frequently occurred and conserved at specific sites in 7 of 8 motifs. Notably, motif **R**XRSXS, R**R**XS, and **R**RXXS with a match of 23, 26 and 20 pArg peptides, respectively, were also the substrate motif of Akt, PKA, and ZIP kinase.^28^ The adjacent pS and pR sites may suggest a potential cross-talk between classical *O*-phosphorylation and non-canonical pArg *N*-phosphorylation, which needs to be investigated by further studies.

### Proglucagon possesses a pArg site located in its vital motif for processing

The paired basic amino acid in pArg motifs, such as RR and KR, is reminiscent of the substrate sequence of propeptide convertase (PC), which functions in the proteolytic activation of proglucagon (PG). Physiologically, peptides cleaved from PG play key roles in blood glucose regulation. This processing is initiated by PC-1 cleaving PG between K^90^R^91^ interdomain and H^92^ residue, resulting in Glicentin (Gli), major PG fragment (MPGF), GLP-1 and GLP-2 (Fig. 4c).^29, 30^ Remarkably, peptide pR^91^HDEFER^97^ derived from PG was repeatedly detected across our MS experiments.

**Figure 4.**
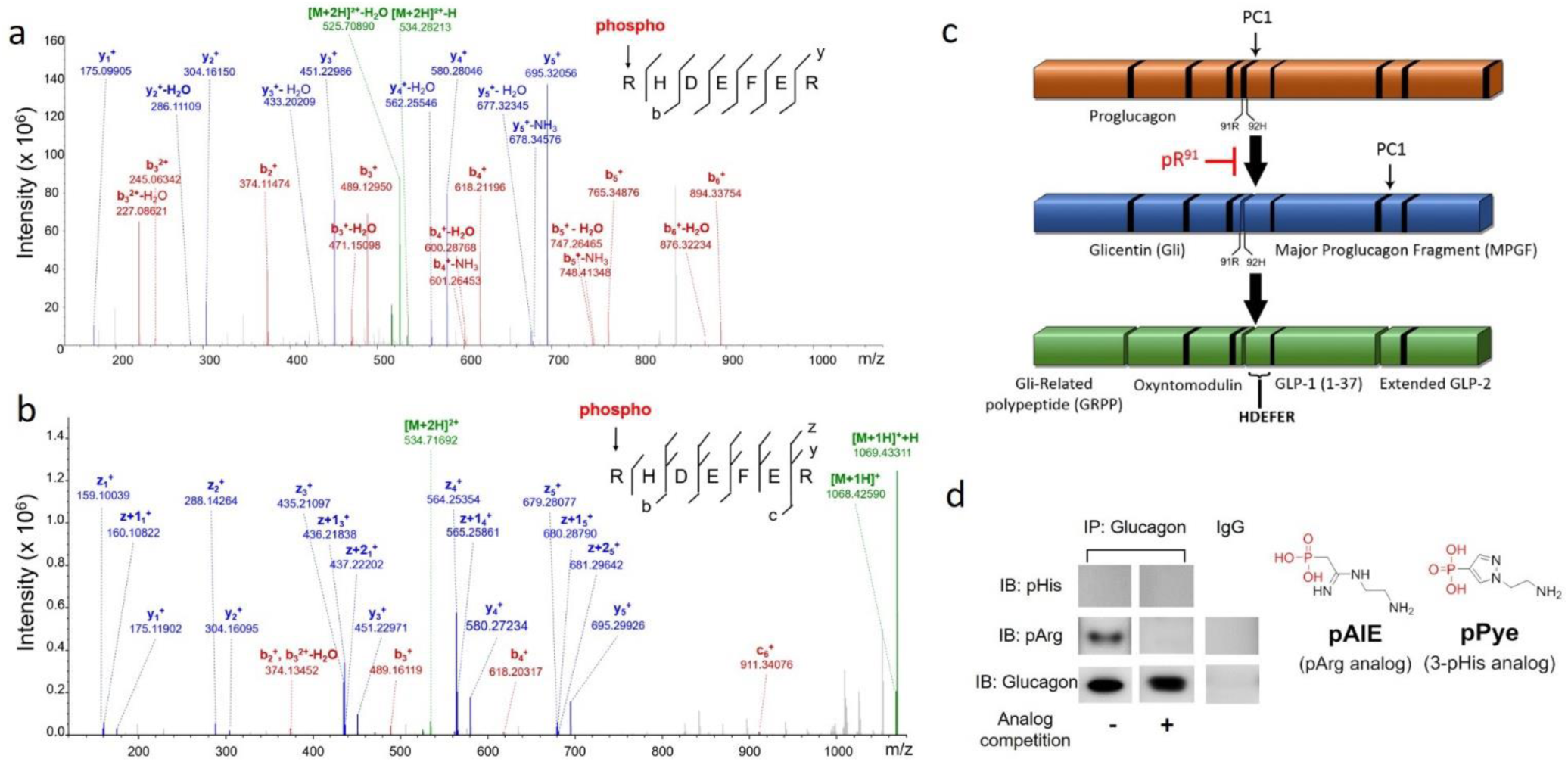
Experimental verification of arginine phosphorylation in proglucagon (PG). **(a)** CID spectrum and (**b**) EThCD spectrum of endogenous PG-derived pRHDEFER peptide from Jurkat cells. All detected fragment ions are summarized in the inset. (**c**) Schematic of the processing of PG by PC1. The first vital cleavage site locates between R^91^ and H^92^. (**d**) Western blotting of PG from Jurkat cell. Immunoprecipitated PG is probed with either anti-pHis or anti-pArg antibody. In the competition assay, the antibody is first incubated with synthesized pPye or pAIE followed by conventional WB.

Since PG have significant biological relevance and K^90^R^91^ interdomain is required for the substrate recognition by PC-1, the site mapping of PG was carefully performed in several ways. As shown in Fig.4a, 4b, both CID and EThcD narrowed down the phosphorylation sites of pR^91^HDEFER^97^ to R^91^ or H^92^, though the conclusive b_1_ ion was missed. The neutral loss of H_3_PO_4_ and HPO_3_ from precursors ([M-80], [M-98]) or b/y-ions, characteristic patterns of O-phosphorylation, were undetectable. Meanwhile, nearly all observed b/y ions exhibited water-loss peaks, indicating a migration of phosphate.^18^ Subsequently, the possibility of phosphorylation on H^92^ was ruled out by comparing the obtained MS^2^ spectrum with that of a synthesized and purified R^91^pHDEFER^97^ peptide (Fig. S13) under the same condition. As shown in Fig. S14 and S15, 1) the y_6_ ion (pH^92^DEFER^97^) assigning to pH^92^, 2) the characteristic pHis neutral loss triplet peaks reported in literature,^31^ 3) pHis diagnostic immonium ion (m/z=190.037) reported recently^16^ were exclusively detectable in the synthetic peptide, but all these characteristics were absent in the cell lysate sample. Biochemically, we immunoprecipitated endogenous PG from Jurkat cell lysate and blotted it with pArg or pHis antibody.^15, 32^ As illustrated in Fig. 4d, no signal was detected by the pHis antibody. In contrast, pArg antibody gave signal, which can be abolished by competition with pAIE, an analog of pArg. Taken together, the phosphorylation site of PG was unambiguously assigned to R^91^ by MS and biochemical validation.

## CONCLUSIONS

Although the widespread occurrence of arginine phosphorylation and its biological relevance have already been elucidated in gram-positive bacteria, revealing it in more complex mammalian cells remains a challenging task. When our manuscript was under revision, the first crystal structure of bacterial pArg kinase McsB is reported 10 years after its discovery.^33^ To boost the study of this unique PTMs, several crucial bottle-necks still need be overcome, that includes 1) Development of solid-phase peptide synthesis for pArg peptides; 2) Generation of sequence-specific pArg antibodies; 3) Identification of mammalian pArg kinase and phosphatase; 4) Determination of pArg stoichiometry in various cell lines and its relationship with other classical *O*-phosphorylation species; 5) Comparison of the pArg levels between physiological and pathological conditions in cells and tissues.

In summary, we reported a new analytical pipeline consisting of antibody-based screening, optimized proteomics workflow, and analog-based competition assay. By virtue of this workflow, we revealed, for the first time, the widespread occurrence of pArg in human cells and its phosphorylation motif. This breaking-through discovery will open a new field for the study of eukaryotic protein post-translational modification. We envision that our analytical workflow together with the identified pArg proteins are expected to be the starting point for both chemical biologists and biochemists towards the studies of mammalian *N*-phosphorylation, a field with enormous challenges and promising prospects.

## METHODS

### Reagents and materials

Conventional reagents such as buffers, inorganic salts, bovine serum albumin (BSA, as blocking agent in WB), phenylmethanesulfonyl fluoride (PMSF), β-mercaptoethanol (BME), acrylamide and bis-acrylamide for electrophoresis, Coomassie brilliant blue G250, N,N,N’,N’-Tetramethylethylenediamine (TEMED) and dithiothreitol (DTT) were purchased from Sangon Biotech (Shanghai, China). cOmplete™ EDTA-free Protease Inhibitor Cocktail and cOmplete™ Protease Inhibitor Cocktail were purchased from Roche Diagnostics (Mannheim, Germany). Acetone used for protein precipitation was purchased from Sinopharm Chemical Reagent (Shanghai, China). LC-MS grade acetonitrile (ACN), formic acid (FA) and ammonium hydroxide (NH_4_OH) were purchased from Sigma-Aldrich (St. Louis, MO, USA). Titansphere TiO_2_ (100Å, 5 µm) was purchased from GL Sciences (Tokyo, Japan). Pre-stained protein ladder (cat. No. 26616) and fetal bovine serum (FBS) were purchased from Thermo Fisher (San Jose, USA). Goat anti-Rabbit IgG HRP conjugate and negative control Rabbit IgG were purchased from ProteinTech Group (Wuhan, China). Amicon ultra centrifugal filter units (MWCO 3500) and PVDF membrane (0.45 µm) were purchased from Millipore (Wisconsin, USA). The sequencing-grade trypsin was from Promega. Potent ECL substrate for western blotting was provided by Smart-Lifesciences (Wuxi, China).

Human embryonic kidney 293T cell line, human lung cancer A549 cell line, human hepatocellular carcinoma HepG2 cell line and human T lymphocyte Jurkat cell line were originally obtained from ATCC and maintained in our laboratory. All cells were cultured in the appropriate medium supplemented with 10 % FBS and 2 mM L-Glutamine according to ATCC’s guidelines. Cell culture media such as DMEM/F-12, DMEM (high glucose), RPMI-1640, and 0.25% trypsin-EDTA were purchased from GE Healthcare Life Sciences (USA).

### Cell culture and pre-treatment

Jurkat cells were cultured in RPMI-1640 (high glucose) at 37 °C, 5% CO_2_ and passaged by direct dilution to fresh medium. Adherent cells were detached with 0.25% trypsin-EDTA at 37 °C and then quenched with complete medium containing 10 % FBS. Cell lines have been tested for mycoplasma contamination annually. Prior to all experiments, cells were allowed to reach ∼70-80% confluence for adherent cells, or a density of 2 × 10^6^ / mL for suspension cells. To elevate the overall phosphorylation level, cells were first washed with warm 1 × PBS, then 1 mM per-vanadate (a pre-mixture of 1 mM Na_3_VO_4_ and 1 mM H_2_O_2_) was added into medium, followed by incubation at 37°C for 30 min before lysis.

### Cell lysis and protein extraction

Adherent cells were detached by trypsinization and then harvested by centrifugation at 800 *g* for 5 min, followed by washing once with cold 1 × PBS. For samples used for western blotting, 5 × 10^6^ cells were lysed with 500 µL of lysis buffer (25 mM Tris, 150 mM NaCl, 1% Triton X-100, pH 8.0) supplemented with 1 mM PMSF, 1 × phosphatase inhibitor cocktails and 1 × protease inhibitor cocktails at 4 °C for 10 min. The lysate was cleared by centrifugation at 14,000 *g* for 10 min at 4 °C. The supernatant was collected and stored at −80 °C for further use. Protein concentration was determined by Bradford method. For samples used for proteomic analysis, the pelleted cells were lysed in a buffer containing 8M urea, 1 mM PMSF, 1 × phosphatase inhibitor cocktails, and 1 × protease inhibitor cocktails at 4 °C for 10 min, and then subjected to the above centrifugation procedures to obtain the supernatant.

### Acid treatment of human cell lysate

5 µL of 1N HCl was added to 20 µL of cell lysate and immediately mixed thoroughly. After 1 h incubating at 30 °C, 5 µL of 1N NaOH was added for neutralization. The protein concentration needs to be < 5 mg/mL; otherwise, proteins were prone to precipitate during this process.

### SDS-PAGE and Western blotting

Cell lysates were diluted with 4 × alkaline loading buffer (160 mM Tris, pH 8.5, 40 % (v/v) glycerol, 4% (w/v) SDS, 0.08 % (w/v) bromophenol blue, and 8 % (v/v) BME) and incubated at room temperature for 10 min without boiling. Proteins were separated on a 10 % SDS-PAGE running in the cold room at 120 V and electro-transferred to PVDF membrane in 0.5 × Towbin buffer containing 20 % (v/v) methanol for 90 min at 4 °C. After blocking with 3 % BSA in TBST (10 mM Tris, pH 8.0, 150 mM NaCl, 0.1% (v/v) Tween-20) for 60 min, the membrane was incubated with the appropriate primary antibody (home-made pan-specific anti-pArg or anti-pHis, 1:1000; anti-proglucagon (Santa Cruz), 1:400) diluted in TBST with 1% BSA for 60 min at RT. Washing step (3 × 5 min) was carried out before and after secondary antibody incubation (1:5000 in TBST). Signals were visualized by incubation with ECL substrate (Smart-Lifesciences, Wuxi, China) for 3 min, and documented with GE Amersham Imager 600 system.

### Synthesis of pPye and pAIE for WB competition assay

Synthesis and characterization of pPye and pAIE were reported by our previous paper,^15^ and they served as antigenic epitope analogs for anti-pHis and anti-pArg antibodies competition assay, respectively. In competitive western blotting assay, the primary antibody was pre-incubated with 2.5 mM of corresponding analogs in TBST buffer (with 1% BSA) at RT for 30 min and then used for standard WB protocol.

### Protein acetone precipitation & tryptic digestion

Jurkat cell extract (∼ 2mg) prepared with Urea-containing lysis buffer, as described above, was subjected to the standard FASP protocol (Wisniewski et al., 2009) or our modified “acetone precipitation protocol for acid-labile proteins”. Briefly, proteins were reduced with 5 mM DTT for 30 min at 25 °C and then alkylated with 10 mM iodoacetamide in the dark for 30 min at 25 °C. Proteins were precipitated by adding 5 volumes of cold acetone and kept at −20 °C for 1 h, and then pelleted by centrifugation at 5000 *g* for 10 min at 4 °C to remove salts and impurities. The pellet was carefully washed twice with cold EtOH and dried in air for 20 min at RT. The pellet was re-suspended in 1 mL of 100 mM NH_4_HCO_3_ (pH 8.0) and sonicated on ice for three cycles (20 sec on, 30 sec off) to homogenize. Sequencing-grade trypsin was subsequently added to the protein solution at a 1:50 ratio (w/w). After 2 h of incubation at 37 °C, another portion of trypsin (1:100, w/w) was added and then incubated for an additional 4 h at 37 °C to ensure complete digestion. The resulting peptide mixture was lyophilized for TiO_2_ enrichment.

### Improved mild-acidic TiO_2_ enrichment

All types of phosphopeptides including acid-stable (pS, pT, and pY) and acid-labile species (pR, pH, and pK) were all enriched from the digested peptide mixture using a modified mild-acidic TiO_2_-based method. Briefly, 6 mg of TiO_2_ was suspended in 1 mL of binding solution (300 mg/mL lactic acid, 12.5% AcOH, 60% ACN, pH 4.0, adjusted with NH_4_OH) and then added to the lyophilized peptide mixture. After gent rotation at 20 °C for 10 min, TiO_2_ was pelleted by centrifugation. The supernatant was collected and incubated with another 1 mg of TiO_2_ for additional 5 min at 20 °C. The TiO_2_ beads from above two steps were combined, and then washed sequentially with 500 μL of binding solution, 500 μL of Wash A (200 mg/ml lactic acid, 75% ACN, pH 4.0), 500 uL of Wash B (200 mg/ml lactic acid, 75% ACN, 10% HAc, pH 4.0) and 500 uL of Wash C (80% ACN, 10% HAc). In each washing step, the TiO_2_ beads were thoroughly re-suspended by pipetting, but the exposure time of the acidic solution was minimized. The enriched phosphopeptides were eluted twice with 100 μL of elution buffer (5% (v/v) of 25 % NH_4_OH in 50% (v/v) ACN) for 15 min each. The eluate was lyophilized and stored at - 80 °C for nano -LC-MS/MS analysis.

### Synthesis of R^91^pHDEFER^97^ peptide

The histidine residue in RHDEFER peptide was chemically phosphorylated with PPA. Briefly, 0.5 mM of RHDEFER peptide was incubated with 50 mM of PPA (pH 8.0) at room temperature for 24 h, and then the resulting RpHEEFER was separated from the unmodified reactant with preparative LC (Solvent A: 0.1 % FA in water, Solvent B: 0.1% FA in ACN).

### Nano-LC-MS/MS analysis

Enriched phosphopeptides were analyzed on Orbitrap Lumos™ Mass Spectrometer (Thermo-Fisher Scientific, USA) equipped with an in-line Easy-nLC 1200 nanoflow LC system (Thermo Fisher Scientific, USA). Lyophilized peptides were dissolved with an appropriate amount of ddH_2_O and immediately loaded from the autosampler onto a homemade C_18_ trapping column (3 µm, 120 Å, SunChrom, USA, 75 µm × 20 mm) at a maximum pressure of 280 bar, then resolved on a homemade C_18_ analytical column (1.9 µm, 120 Å, 150 µm × 150 mm) with a gradient of 95% A (0.1% FA), 5% B (0.1% (v/v) FA in 80% ACN) to 65% A, 40% B over 75 min at 600 nL/min. The effluent was introduced via a nano-ESI source with 2 kV capillary voltage. Survey scans were acquired over 380-1,400 m/z at a resolution of 120K in DDA approach in positive mode. The ten most abundant ions were selected for fragmentation by CID or CID-triggered EThcD or EThcD in the top-speed mode with a 60-sec dynamic exclusion. The precursor automatic gain control (AGC) target was set to 4 × 10^5^, and the maximum injection time was set to 50 ms. For CID, 35 % normalized collision energy (NCE) was applied. For EThcD, the reaction time was set to 100 ms (AGC target: 4 × 10^5^, maximum injection time: 100 ms), and HCD NCE was set to 30%. For CID-triggered EThcD, 25 % NCE was applied for CID, and the precursors with NL (m/z = 79.97 or 97.9763) were selected for EThcD fragmentation (AGC target: 4 × 10^5^; maximum injection time: 150 ms; ETD: Calibrated Charge-Dependent parameters; HCD CE: 25%).

### Proteomics data analysis

Raw data files were processed using Proteome Discoverer Software (version 2.1.0.80, Thermo-Fisher Scientific) and searched against a combined forward/reversed database of *homo sapiens* UniProt Reference Proteome (2015.12.02; 20,187 sequences) and common contaminants (10,540 entries in total). Mascot and SEQUEST (version 27, Thermo Fisher Scientific) algorithm were used with the following parameter settings (Precursor mass error tolerance: 10 ppm; Fragment mass error tolerance: 0.5 Da; Enzyme: trypsin; Max. missed cleavages: 3; Fixed modifications: carbamidomethylation (C); Variable modifications: acetyl (protein N-terminus), oxidation (M), phosphorylation (S, T, Y, H, K, R); Max. variable PTM per peptide: 4; Fragment ion: *b* and *y* ions (for CID) and *b, y, c* and *z* ions (for EThcD); Target FDR for proteins, peptides and PSMs: 0.01) Phosphopeptide hits were analyzed using *ptm*RS in phosphoRS node. All data files were exported to EXCEL files for statistical analysis.

### Motif analysis

The phospho-sites (pR, pH and pK) identified with *ptm*RS ≥ 75% and the surrounding residues (+/- 6) were analyzed by the Motif-X algorithm (http://motif-x.med.harvard.edu/). Human IPI database was used as a background. Sites that can’t be extended due to the N- or C-terminus were excluded. The significance threshold (*p*-value) was set to < 10^-6^, and the minimum number of motif occurrences was set to 20.

### Data availability

The MS proteomics data were deposited to the ProteomeXchange Consortium (http://proteomecentral.proteomexchange.org) via the PRIDE^34^ partner repository with the dataset identifier PXD009696. All other data supporting the findings of this study are available from the corresponding author on reasonable request.

## ASSOCIATED CONTENT

Supporting Information

The Supporting Information is available free of charge on the ACS Publications website at DOI:

List of all identified peptides including phosphopeptides (STYRHK) and non-phosphopeptides before and after per-vanadate treatment of Jurkat cell (EXCEL 1); List of high-confident *N*-phosphosites and proteins (RHK) used for phosphorylation motif and GO analysis (EXCEL 2); Gene Ontology (GO) analysis of high-confident *N*-phosphorylated proteins (RHK) (EXCEL 3);

## Author Contributions

C. Fu conceived this study and wrote the manuscript; SS. Fu, Q. Zhuo, RC. Lin, H. Ouyang, MN. Wang and Y. Sun performed the experiments; SS. Fu and C. Fu carried out data analysis.

## Notes

The authors declare no competing financial interest.

## ACKNOWLEDGEMENT

We thank Dr. J. Qin and Dr. MW. Liu (The PHOENIX Center, Beijing) for helpful discussion, Dr. R. Ding and YH. He (Xiamen University) for help in data analysis, Dr. Andrew Ferenbach and Mr. Andrii Gorelik (University of Dundee) for proofreading the manuscript.

